# Modified E2 glycoprotein of hepatitis C virus enhances pro-inflammatory cytokines and protective immune response in mice

**DOI:** 10.1101/2022.02.07.479452

**Authors:** Vijayamahantesh, Tapas Patra, Keith Meyer, Alameh Mohamad-Gabriel, Erin Reagan, Drew Weissman, Ranjit Ray

## Abstract

Hepatitis C virus (HCV) is characterized by a high number of chronic cases owing to an impairment of innate and adaptive immune responses. CD81 on the cell surface facilitates HCV entry by interacting with the E2 envelope glycoprotein. On the other hand, CD81/E2 binding on immune related cells may also influence host response outcome to HCV infection. Here, we performed site-specific amino acid substitution in the front layer of E2 sequence to reduce CD81 binding and evaluate HCV candidate vaccine potential. The altered sE2 protein (F442NYT), unlike sE2, displayed a significant reduction in CD81 binding, induced pro-inflammatory cytokines, and repressed anti-inflammatory response in primary monocyte-derived macrophages as antigen presenting cells. Further, sE2_F442NYT_ stimulated CD4^+^T cell proliferation. Immunization of Balb/c mice with an E1/sE2_F442NYT_ RNA-lipid nanoparticle (LNP) displayed improved IgG1 to IgG2a isotype switching, an increase in HCV pseudotype virus neutralizing antibodies, and resistance to challenge infection with a surrogate recombinant vaccinia virus expressing HCV E1-E2-NS2_(aa134-966)_, unlike parental E1/sE2 immunization. Further investigation on modified E2 antigen for selection as antigen may provide helpful information for HCV vaccine development.

**One Sentence Summary:** Reduced HCV E2-CD81 binding and immune response

## INTRODUCTION

HCV causes chronic liver disease in many infected humans and is one of the contributing factors for cirrhosis and hepatocellular carcinoma. Direct-acting antiviral agents (DAAs) have transformed the standard therapy, achieving high rates of sustained virologic response (SVR). Unfortunately, despite anti-viral efficacy, DAAs cannot fully eradicate liver cancer risk, especially after HCV clearance with advanced liver disease (*1, 2*).

HCV has evolved mechanisms to evade immune activation and resolution of infection. Several studies of acute HCV infection demonstrate that virus clearance is associated with broad and potent T-cell activation (*3-5*), and a rapid induction of cross-reactive neutralizing antibody responses (*6-8*). A Phase I vaccine trial of recombinant HCV E1/E2 envelope glycoproteins in healthy human volunteers was conducted (*9-11*). IL-10 secretion by PBMCs was observed, and generation of a variable range of IL-4 secretion was noted with the candidate HCV E1/E2 vaccine. Vaccination failed to exhibit a clear dose dependent response to escalated E1/E2 protein administration, and only ∼28% of the vaccinated human sera displayed a detectable virus neutralization response. A recent report suggests that vaccination of human subjects with a recombinant virus vector expressing non-structural proteins lowered HCV RNA level, but did not prevent chronic infection (*12*).

We have subsequently shown that HCV E2 induces the immune regulatory cytokine; IL-10, and CD163 protein from primary macrophages (*13*). Further, HCV E2 enhances STAT3 and suppresses STAT1 activation, suggesting macrophage polarization towards the M2 phenotype. HCV E2 interrupts the function of T-, B- and NK- cells by binding with CD81 (*14-17*). HCV-specific CD4^+^T-cell immune deficiency may be the primary cause of CD8^+^T-cell functional exhaustion (*18, 19*).

HCV E2 contains multiple epitopes for both T and B cells (*20-22*). We examined whether impairing the E2-CD81 interaction can promote T-helper cell functions to induce a robust HCV E2 antigen specific immune response. Our results suggest that reduced level HCV E2-CD81 interaction improved immune response, contributing to the development of an HCV vaccination strategy.

## RESULTS

### Amino acid mutation in E2 front layer impairs CD81 binding and alters cytokine response

The front layer of the HCV E2 glycoprotein interacts with the large extracellular loop (LEL) of CD81 in the process of viral entry (*23, 24*). The front layer is located within two hypervariable regions of the HCV E2 ectodomain. We have postulated two different approaches from an earlier report (*25*) on structural and CD81 binding analyses of E2 glycoprotein to design variants to disrupt CD81 interaction. For one such approach, we selected one amino acid substitution by replacing non-polar leucine to polar tyrosine at position 427 (L427Y). In another approach, we introduced a N-linked glycosylation site (N-X-T/S) at positions 442 and 444 on the front layer of E2 (F442NYT) to abrogate E2-CD81 binding with a hyper-glycosylated form (Fig. 1, panel A). The effect of impaired E2-CD81 binding on the immunogenicity of HCV E2 was investigated by performing the amino acid substitution for generation of E2 variants from a sE2 lacking the hydrophobic C-terminal transmembrane (384-661) of HCV genotype 1a H77 strain (Fig. 1, panel A). sE2 or modified glycoproteins were expressed in mammalian HEK293T cells, known to attach different N-linked glycosylation moieties to proteins. Secreted sE2 and other variant proteins bearing a C-terminal hexa-histidine tag were purified from culture supernatants by immobilized nickel affinity chromatography for subsequent characterization. ELISA was performed to test E2-CD81 interaction of different variants of secretory E2 collected from the culture fluid of transfected constructs, and the purified proteins. A significant reduction in CD81 binding for sE2_L427Y_ (∼60% loss) and sE2_F442NYT_ (∼70% loss) as compared to the sE2 protein available from culture fluid. Purified protein from sE2_L427Y_ (∼35%) and sE2_F442NYT_ (∼40% loss) also exhibited a difference in binding activity. (Fig. 1, panels B and C). Thus, amino acid substitutions in the front layer of CD81 binding domain of E2 resulted in impairment of CD81 interaction. We continued further study utilizing E2_F442NYT_ construct for understanding immunoregulatory activity and evaluated as a candidate vaccine antigen for protective response.

**Fig. 1.**
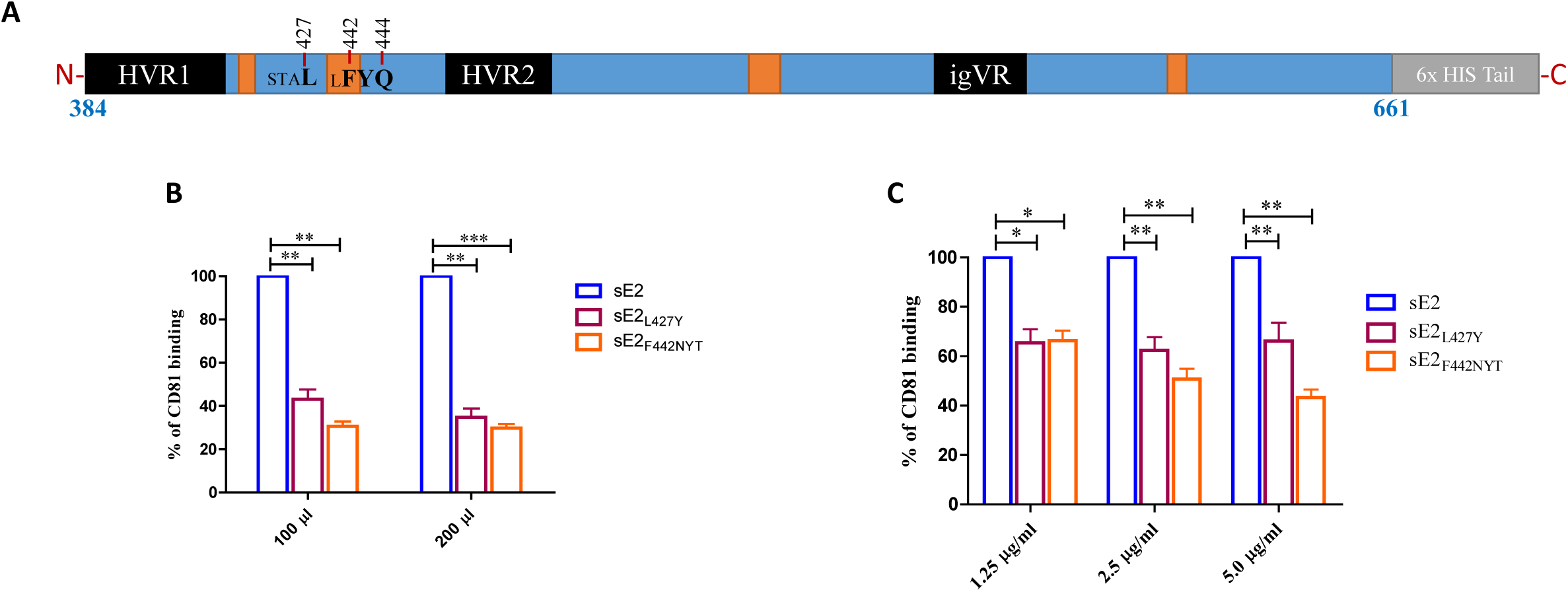
Schematic presentation of HCV sE2 ectodomain and the mutants. Amino acid positions of interest for mutations in a linear map of sE2 are indicated (panel A). Comparison of human CD81 binding with modified E2 expressed transiently in HEK293T transfected cell culture supernatant with increasing volumes (panel B) and purified histidine tag recombinant proteins in a dose dependent concentration (panel C) are shown. Culture medium from mock transfected HEK293T cells or PBS were used as reagent control. The significance levels are expressed as *=p<0.05 ** = p < 0.005, *** = p < 0.001.

### Inhibition of E2/CD81 skews IL-10/IL-12 response in human macrophage cell line

Both macrophages and DCs play an important role in immune surveillance. We have previously shown that HCV E2 significantly inhibits macrophage polarization toward the M1 phenotype, and antigen presenting DC maturation (*11, 13, 26*). HCV E2 binds to its cognate receptor; CD81, on macrophages and induces the production of the immune suppressive cytokine, IL-10. We hypothesized that a loss of E2/CD81 binding on macrophages would reduce the production of IL-10 by macrophages. Initially, we tested our hypothesis in a human monocyte cell line, THP-1. We stimulated THP-1 in the presence or absence of HCV sE2, sE2_L427Y_, or sE2_F442NYT_ *in vitro*. A significantly increased production of IL-10 from THP-1 by sE2 compared to modified sE2 proteins was observed (Fig. 2, panel A). In contrast, altered sE2 proteins induced an increased IL-12 production compared to sE2 and unstimulated control (Fig. 2, panel B). The present observation strengthens our hypothesis that binding of sE2 to CD81 results in the production of IL-10 and reduced binding inhibits IL-10 production.

**Fig. 2.**
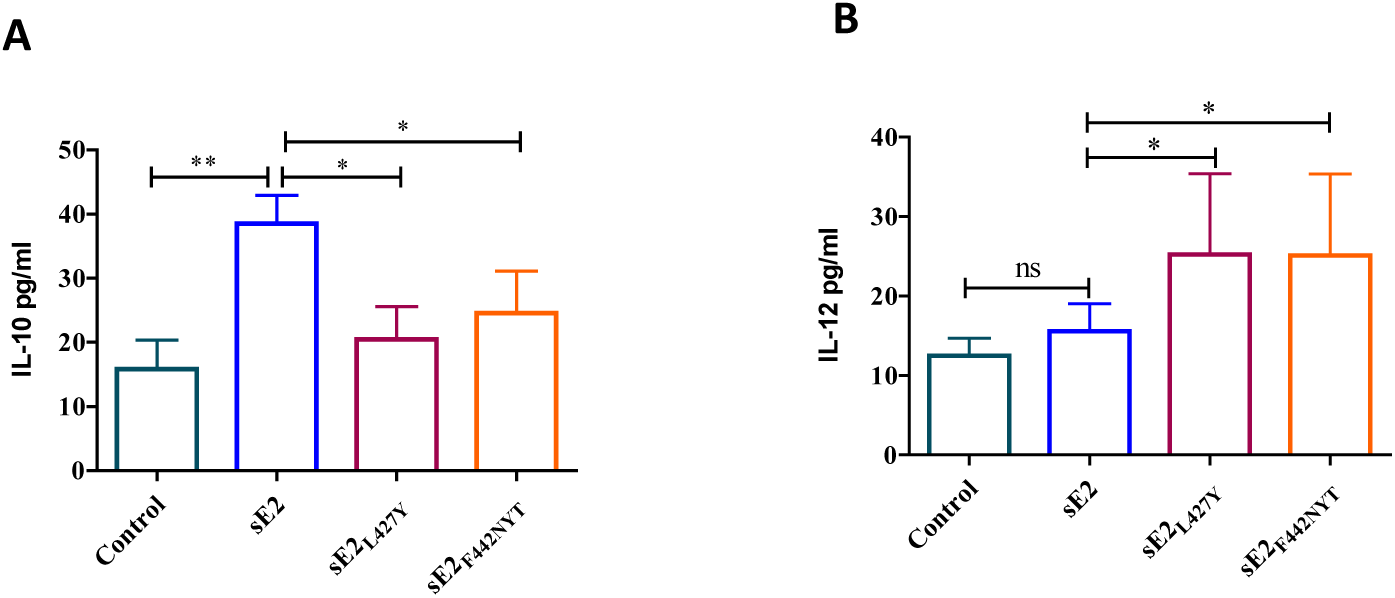
IL-10 and IL-12 cytokine expression status following incubation of THP-1 cells with modified E2 proteins. Human monocyte derived macrophage cell line THP-1 was treated with 2.5 µg/ml of modified E2 for 24 h. Secretory cytokines IL-10 and IL-12 in the culture supernatant were quantified by ELISA (panels A and B). The significance level is expressed as *=p<0.05 **= p < 0.005.

### Inhibition of E2/CD81 binding induces pro-inflammatory cytokine production

We analyzed immune stimulatory effect of the modified E2 on primary human monocyte derived macrophages. Secretion of pro-inflammatory cytokines was measured in the culture supernatant of modified E2 exposed macrophages. sE2 induced significantly higher IL-10 generation from macrophages as compared to the altered sE2 (sE2_L427Y_, or sE2_F442NYT_) proteins (Fig. 3, panel A). Within the altered sE2 group, the sE2_L427Y_ generated more IL-10 as compared to sE2_F442NYT._ The presence of IL-10 has an inhibitory effect on the induction of IFN-γ, and associated genes (*27*). In contrast to IL-10 generation, we observed a significant induction of pro-inflammatory cytokines from modified sE2 as compared to sE2 and unstimulated control macrophages. IL-4 induces differentiation of naïve T cells to the Th2 phenotype. As seen with IL-10, a significant increase in the production of IL-4 was apparent in macrophages incubated with the sE2 protein, and this effect was mitigated in the altered sE2 proteins (Fig. 3, panel B). Among the tested pro-inflammatory cytokines, IL-12 induction was consistently higher in macrophages exposed to modified sE2_L427Y_, or sE2_F442NYT_ proteins (Fig. 3, panel C). Exposure of macrophages to sE2 generated little production of IFN-γ above control (Fig. 3, panel D). On the other hand, a significant increase in IFN-γ levels was measured from macrophages exposed to sE2_F442NYT_ protein. Within the treatment groups, sE2_F442NYT_ produced significantly higher IL-6 compared to sE2_L427Y_ and sE2 (Fig. 3, panel E). We also observed a significantly higher TNF-α generation from sE2_L427Y_, or sE2_F442NYT_ protein (Fig. 3, panel F). The pro-inflammatory vs anti-inflammatory cytokine ratio increased with inhibition of E2/CD81 interactions.

**Fig. 3.**
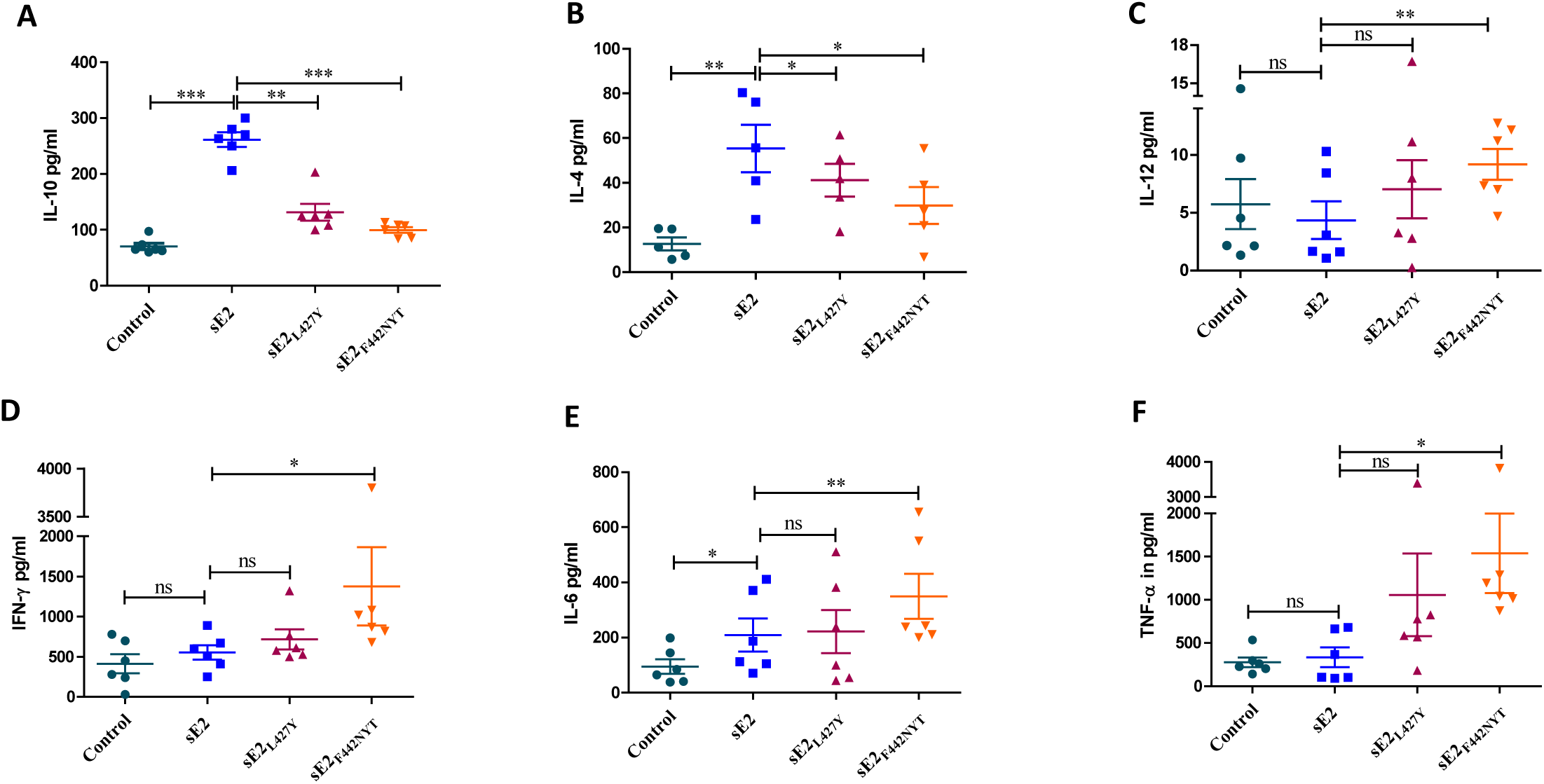
Cytokine expression from macrophages of healthy donors incubated with sE2 or modified E2 for comparison. Monocyte derived macrophages from healthy donors (n=6) were incubated with 2.5 µg/ml HCV sE2 or modified proteins for 24 h, and secretory cytokines (IL-10, IL-4, IL-12, IFN-γ, IL-6, and TNF-α) in the culture supernatant were quantified by ELISA (panels A-F). The significance level is expressed as *=p<0.05, ** = p < 0.005, *** = p < 0.001, or ns (not significant).

### sE2_F442NYT_ induced CD4^+^T cell proliferation

Proliferative CD4^+^T cell response against HCV proteins is a strong immunological correlate of the outcome of acute HCV infection (*28*). Proliferative HCV-specific CD4^+^T cell responses are usually not detectable in acute persisting and chronic HCV infection. Here, we analyzed the role of modified E2 in promoting isolated CD4^+^T cell proliferation. Cells isolated from healthy volunteers were incubated with purified sE2 or sE2_F442NYT_ for 72hrs in the presence of CFSE. FACS analyses revealed a significant impairment in CD4^+^T cell proliferation following sE2 treatment (Fig. 4). In contrast, sE2_F442NYT_ treated cells displayed similar proliferation response as control. Thus, the results suggest that sE2_F442NYT_ mutant does not inhibit CD4^+^T cell proliferation unlike sE2.

**Fig. 4:**
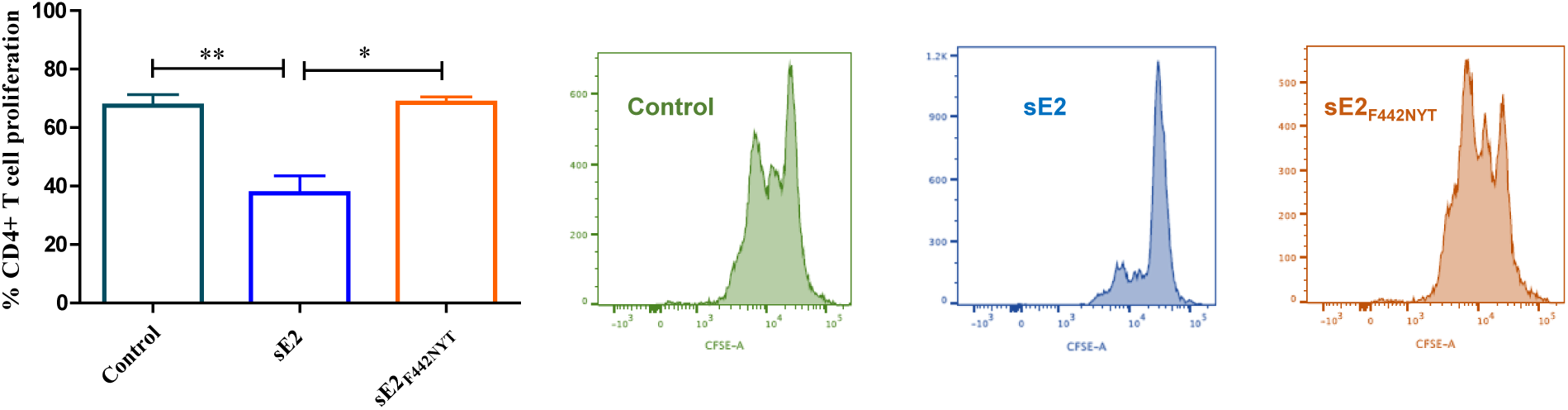
sE2_F442NYT_ induced CD4^+^T cell proliferation. CD4^+^T **c**ells isolated from healthy volunteers were incubated with purified sE2 or sE2_F442NYT_ for 72hrs in the presence of CFSE. Proliferation results from FACS analyses are shown. The significance levels are expressed as *=p<0.05 ** = p < 0.005.

### sE2_F442NYT_ activated macrophages promote CD4^+^T cell differentiation into Th1 type

Differentiation of naive T cells into Th1 effector type is critically required to induce a protective immune response against intracellular pathogens (*29*). Polarization of Th0 into Th1 or Th2 type effectors depends on the type and concentration of cytokines present at the immune synapse, the strength of T cell receptor-mediated signals, and the activation state of the antigen presenting cells. As sE2_F442NYT_ activates the macrophages towards M1 type, and induces the production of pro-inflammatory cytokines, we hypothesized that these activated macrophages could polarize T cells towards Th1 type effector cells. To test our hypothesis, we incubated macrophages in the presence or absence of sE2 or sE2_F442NYT_ for 24 hours and co-cultured with autologous CD4^+^T cells. CD69 surface expression correlates with differentiation of CD4^+^T cells into Th1 effector cells (*30*). We observed an increased expression of the CD69 activation marker on both sE2 and sE2_F442NYT_ treated as compared to untreated CD4^+^T cells. However, expression of CD69 on sE2_F442NYT_ treated T cells was significantly (p<0.005) higher as compared to sE2 treated T cells (Fig. 5, panel A). We did not detect a significant change in surface CD25 expression in cells co-cultured with sE2 or sE2_F442NYT_ treated macrophages after 24 hours (Fig. 5, panel B). We observed a significant increase in the Th1 marker; CXCR3, on CD4^+^T cells co-cultured with sE2_F442NYT_ compared to untreated or sE2 treated macrophages (Fig. 5, panel C). However, we could not find any significant difference (p>0.05) in the expression of the Th2 marker CCR4 on CD4^+^T cells between the different treatment groups (Fig. 5, panel D).

**Fig. 5:**
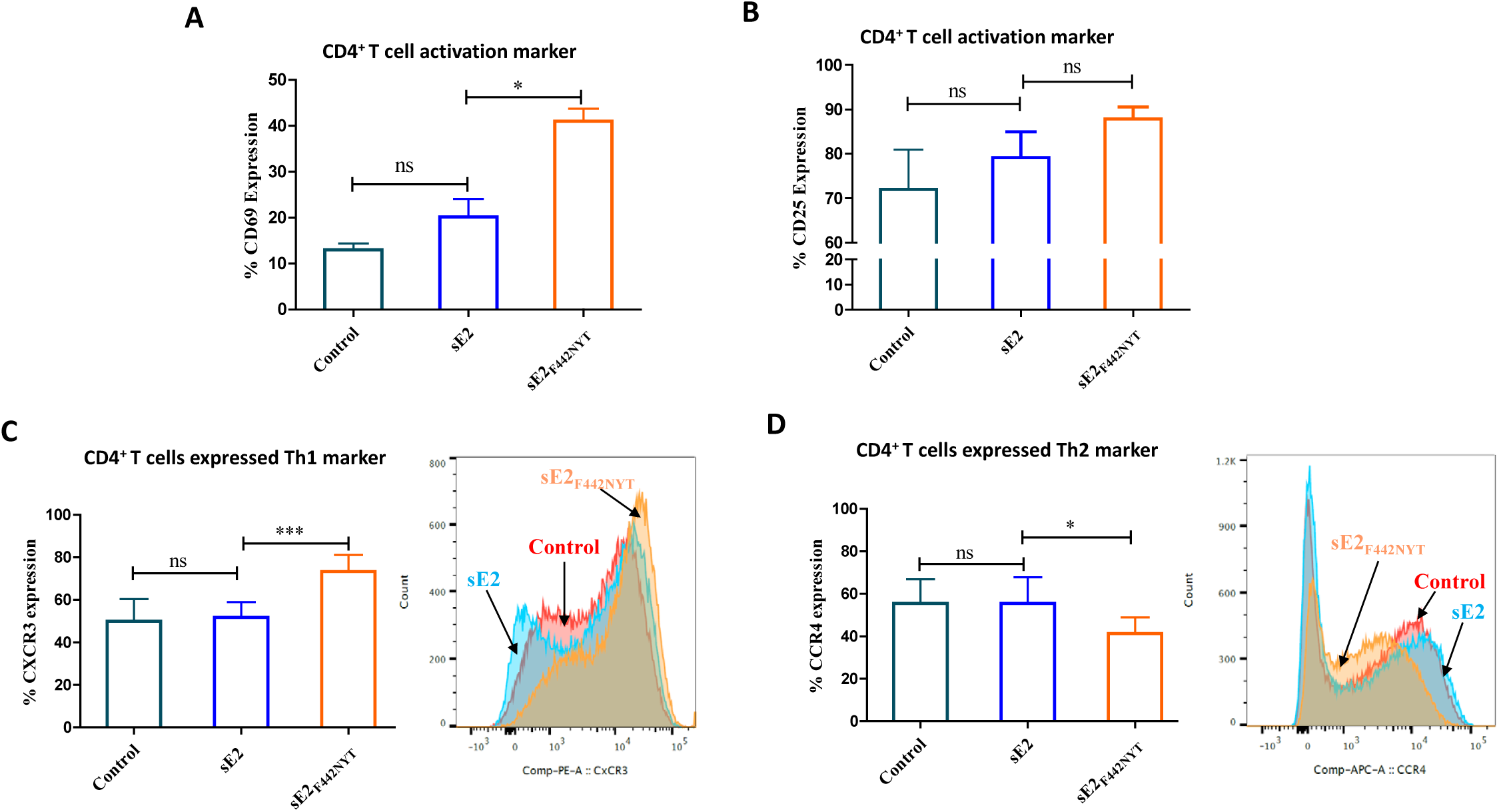
The effect of macrophage activation by sE2_F442NYT_ on polarization of CD4^+^T cells toward a Th1 type. CD4^+^T cells were isolated from healthy donors (n=4) and treated with sE2 or sE2_F442NYT_ for 2 hrs. in the presence of human anti-CD3 (5µg/ml) and anti-CD28 (1µg/ml). The expression of T cell activation markers CD69 and CD25 on the surface of CD4^+^T cells were quantified by Flowcytometry (panels A and B). Monocyte derived macrophages isolated from healthy donors (n=6) were treated with sE2 or sE2_F442NYT_ protein for 24 hrs and autologous CD4^+^T cells were co-cultured with the treated macrophages for 4 days. Th1 and Th2 polarization were measured by expression of CXCR3 and CCR4, respectively, on the surface of CD4^+^T cells by flowcytometry (panels C and D). The level of significance is expressed as *=p<0.05, *** = p < 0.001, and ns (not significant).

### Immunization of mice with E1/sE2_F442NYT_ mRNA-LNP vaccine induces pro-inflammatory cytokines

We tested the immune stimulatory effect of the modified sE2 in mice immunized with E1/sE2 or E1/sE2_F442NYT_. E1/sE2 construct induced significantly higher IL-10 production in serum compared to that of the E1/sE2_F442NYT_ vaccine preparation and was highly significant compared to unstimulated control (<0.01) (Fig. 6, panel A). This observation supports our previous report that the ablation of E2/CD81 interaction on macrophages reduces E2 induced IL-10 production. IL-10 has an inhibitory effect on IFN-γ production and suppression of IFN-γ induced genes (*27*). We observed a significant induction of IFN-γ in contrast to IL-10 production in the E1/sE2_F442NYT_ vaccinated mice as compared to sE2 and unstimulated control macrophages (Fig. 6, panel B). Among the tested pro-inflammatory cytokines, IL-12 production was enhanced in the E1/sE2_F442NYT_ immunized mice (Fig. 6, panel C). Within treatment groups, E1/sE2_F442NYT_ produced significantly less IL-4 as compared to E1/sE2 (Fig. 6, panel D). Taken together, immunization of mice with E1/sE2_F442NYT_ mRNA vaccine reversed the IL-10/IL-12 secretion ratio, and led to enhanced serum IFN-γ, reducing IL-4 levels.

**Fig. 6:**
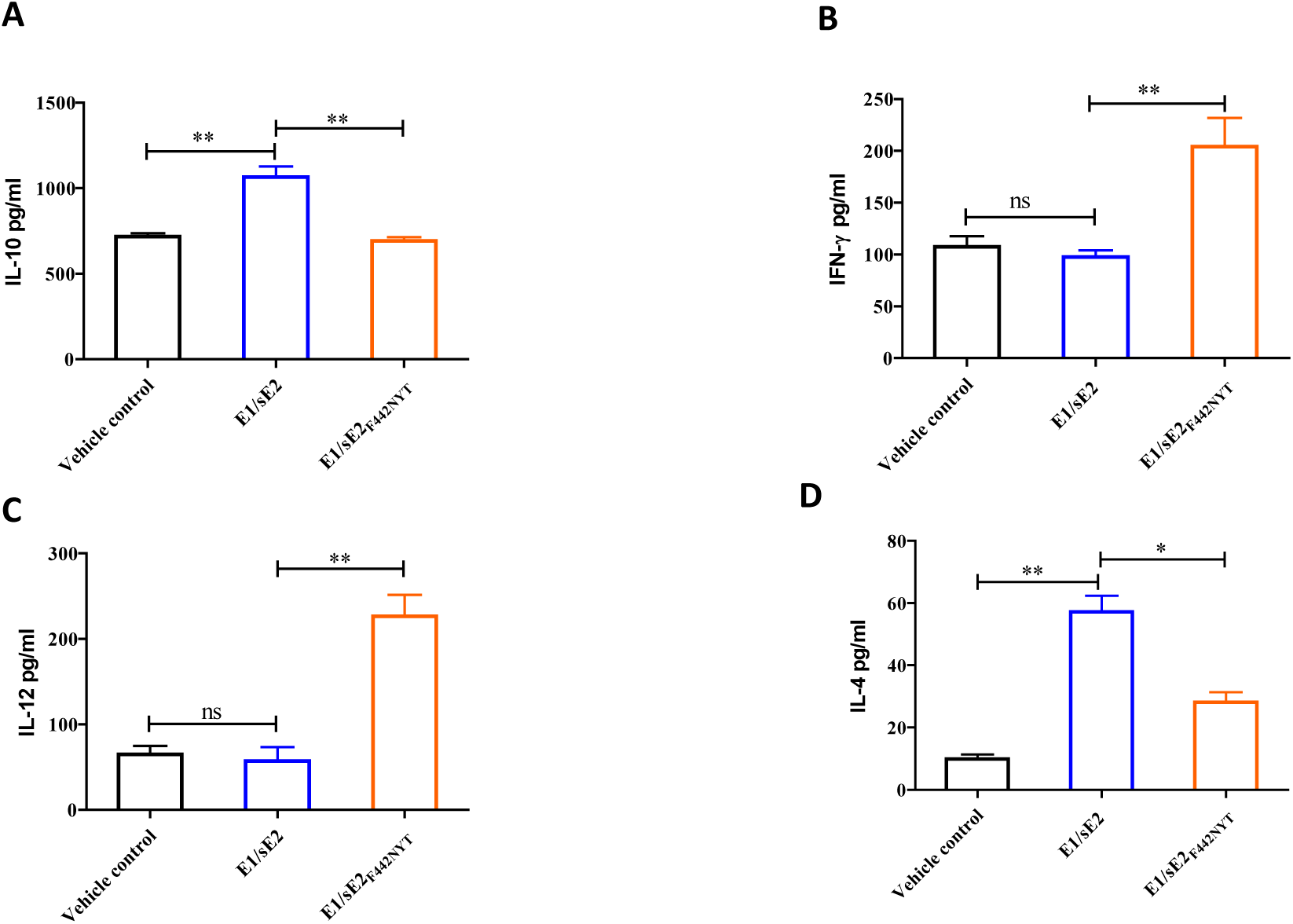
Comparison of mouse cytokine response following immunization with E1/sE2 or E1/sE2_F442NYT_. IL-4, IL-10, IL-12 and IFN-γ expression (panels A-D) were quantified from sera of unimmunized control, E1/sE2, or E1/sE2_F442NYT_ vaccinated mice by ELISA. The significance level is expressed as *=p<0.05 ** = p < 0.005, or ns (not significant).

### Qualitative nature of antibody improved from E1/sE2_F442NYT_-mRNA vaccine in mice

Antibody isotype switching occurs post antigen exposure from IgM to other Ig isotypes. The Th1 subtype of CD4^+^T cells secrete IFN-γ, which induces an isotype switch in B cells to IgG2a secretion. Whereas Th2 cells are associated with enhanced IgG1 production. Here, mice immunized with the E1/sE2-mRNA vaccine preparation exhibited pronounced IgG1 in serum. In contrast, a distinct skew in the isotype of the E1/sE2_F442NYT_ mRNA immunized mice was apparent towards IgG2a production (Fig. 7, panel A). The HCV-lentiviral pseudotype particle (HCVpp) system was used to analyze relative neutralization activity between these two immunogen preparations. Serum recovered from mice immunized with the E1/sE2_F442NYT_-mRNA preparation displayed a minimum of a 2-fold enhancement in neutralization efficacy on HCVpp, as compared to the E1/sE2 immunized mice (Fig. 7, panel B).

**Fig. 7:**
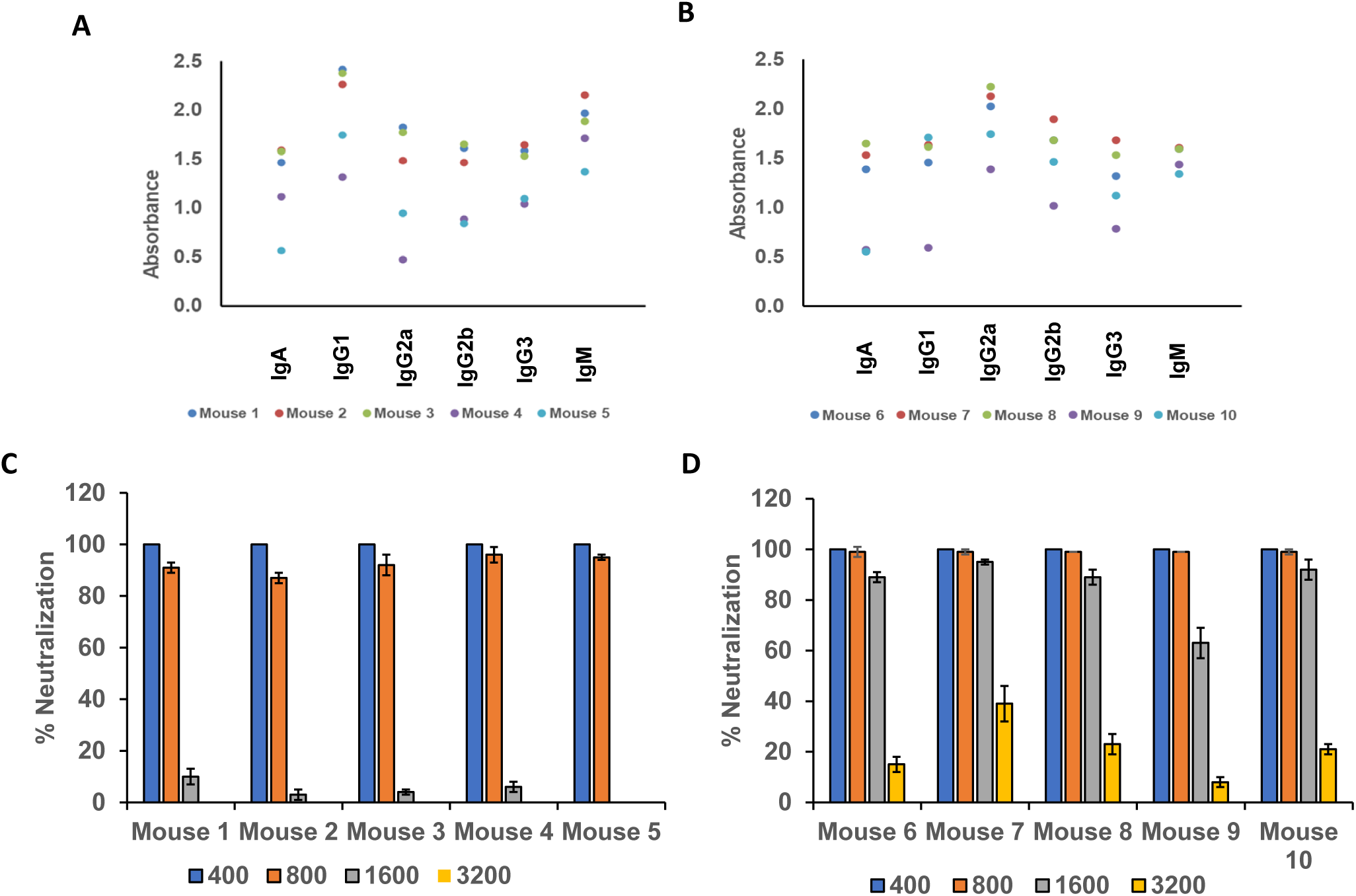
Mouse isotype specific antibody response following immunization with E1/sE2 or E1/sE2_F442NYT_-mRNA-LNP. Immunized mouse sera (collected after 4 weeks) were analyzed for isotype specific antibody response by coating purified E2 antigen on commercially available ELISA plate for isotype antibody analyses. Mouse 1-5 denotes E1/sE2 and mouse 6-10 denote E1/sE2_F442NYT_-mRNA-LNP immunized animal sera (panels A and B). Neutralization of HCVpp as a surrogate virus was carried out using individual sera at different dilutions (1/400 – 1/3,200) from the same immunized animals (panels C and D).

### E1/sE2_F442NYT_ mRNA-LNP vaccine enhances protective immune response

E1-sE2_F442NYT_ mRNA-LNP immunized mice were sacrificed after 4 days of challenge infection with surrogate recombinant vaccinia virus expressing HCV E1-E2-NS2_aa134-966_. Ovary from experimental animals were homogenized in 300 ul medium, freeze-thawed 3 times and centrifuged to pellet cell debris. Clear supernatant was serially 10-fold diluted and plaque assayed on a BSC40 cell monolayer. Plaques were stained after 3 days with 1% crystal violet and counted. Interestingly, vehicle control or wild type E1/sE2 mRNA vaccinated mice failed to show a statistically different level in virus recovery by plaque assay (Fig. 8). In contrast, the E1/sE2_F442NYT_ mRNA vaccine displayed a significant reduction (≥ 4 log) in virus recovery from the ovary of challenged mice. Two out of 5 mice showed minimal virus recovery, while the remaining 3 displayed an undetected virus titer (Fig. 8). In conclusion, the E1/sE2_F442NYT_ mRNA-LNP vaccine preparation induced a significantly improved protective response in immunized animals.

**Fig. 8:**
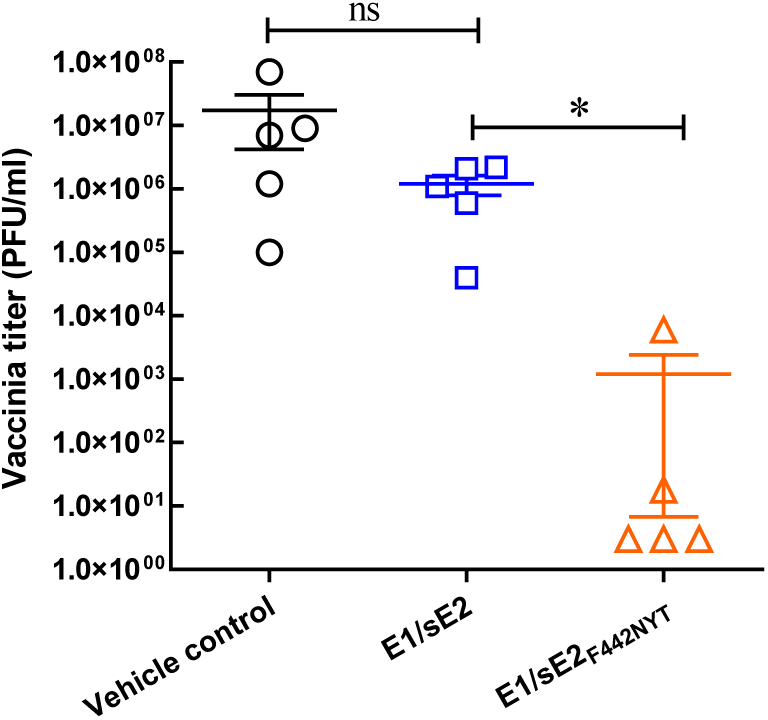
Protection of immunized mouse following challenge infection of recombinant VV expressing HCV E1-E2-NS2_aa134-966_. Female Balb/c mie (5 in each group) were immunized with vehicle control, E1/sE2, or E1/sE2_F442NYT_-mRNA-LNP. Mouse ovaries were collected after rVV challenge infection, homogenized in medium, freeze-thawed 3 times, and centrifuged to pellet cell debris. Clear supernatant was serially 10-fold diluted and plaque assayed for virus recovery on BSC40 cells. Crystal violet-stained plaques were counted, and expressed as pfu/ml. The level of significance is expressed as *=p<0.05 or ns (not significant).

## DISCUSSION

HCV E2 induces the immune regulatory cytokines and interrupts the function of immune related cells by binding with CD81 (*9-10, 13-17*). Here, we examined whether the soluble modified E2 antigen induces an improved immunogenic profile for vaccine development. We observed a gain in CD4^+^T helper cell function by modulated E2 for loss of CD81 binding activity and inclusion of a potential glycosylation site in a mRNA-LNP vaccine prepared of E1/sE2_F44NYT_ mutant. Protective HCV immunity in mice was noted upon challenge infection with a surrogate recombinant vaccinia virus. Previously, purified HCV E1/E2 envelope glycoproteins together with MF59 adjuvant in a safety and immunogenicity vaccine study in humans suggested induction of a weak immune response (*9*). MF59 was used in the vaccine trial to facilitate a Th1 cytokine profile and induction of CD4^+^ memory and cytotoxic T-lymphocyte responses. However, the vaccinated volunteers displayed induction of IL-10, with minimal increase in IL-12 expression. Our *in vitro* study suggested that HCV E2 induced IL-10 in human PBMC derived macrophages, with minimal IL-12 production (*13*), and M2 phenotype polarization of macrophages. Macrophages play a crucial role in antigen presentation, and in the interaction between innate and adaptive immunity. M1 macrophages promote a Th1 response and possess antiviral activity, while M2 macrophages are involved in the promotion of a Th2 response, and immune tolerance (*31, 32*).

Glycosylation has a role in adaptive immune activation (*33*). Appropriate glycosylation of antigens is exploited for the development of more efficacious prophylactic as well as therapeutic vaccines (*34*). E2 front layer constitutes most of the CD81 binding sites and is remarkably flexible (*35*). The E2 front layer (residues 424-453) is suggested to be highly immunogenic skewing the immune response to non-neutralizing epitopes during immunization. E2 is densely glycosylated and the CD81 receptor-binding site is not obstructed by any of the 11 N-linked glycans (25). Removal of N-linked glycosylation sites does not appear to alter antigenic properties of HCV E2 (*36*). Our results from the present study indicated that interruption of CD81 binding and introduction of a potential N-linked glycosylation site on the modified sE2 antigen improved the way for gain in T cell response that switched the generation of anti- to pro-inflammatory cytokines compared to single amino acid substitution in sE2_L427Y_. The modified sE2_F442NYT_ helped in the generation of pro-inflammatory cytokines (IFN-γ, IL-12, IL-6 and TNF-α) while sE2_L427Y_ did not to such an extent. Thus, the presence of a potential glycosylation site on sE2_F442NYT_ appeared to help in better antigen presentation to T lymphocytes for immune activation.

Disrupting CD81 binding along with the inclusion of a potential glycosylation site as sE2_F442NYT_ surprisingly contributed a stronger cytokine profile change, unlike the loss of CD81 binding in sE2_L427Y_. We observed that p-Stat3(S705) is not induced by sE2_F442NYT_, unlike sE2 or E2_L427Y,_ although Stat1 was activated by either of the E2 mutants, implying decreased IL-10 expression. Stat3 activation may be related for dampening cytokine mediated immunoregulatory activity by HCV E2 as suggested earlier (13).

Cytokines play an important role in the homeostasis of the immune system, attributing to the development of subsequent resistance or susceptibility to disease (*37, 38*). IL-10 is a pleiotropic master regulatory cytokine produced by various immune cells; primarily macrophages, dendritic cells and T-helper cells. IL-10 critically associates with the persistence of viruses by suppressing cell mediated immunity and represents a major immunosuppressive cytokine. The role of IL-10 is to limit the extent of the activation of both innate and adaptive immune cells to maintain a homeostatic state. IL-10 regulates growth and/or differentiation of B cells, NK cells, cytotoxic and helper T cells, mast cells, granulocytes, dendritic cells, keratinocytes, and endothelial cells (*39*). On the other hand, IL-12 orchestrates resistance against infectious diseases through macrophage activation, and induction of IFN-γ, and helps in eliminating intracellular pathogens (*40*). IL-12 and IFN-γ work concomitantly in the development of a Th1 immune response against viral infections. Exposure of macrophages to IL-10 suppresses IFN-γ induced genes and prevents them from responding to IFN-γ (*27, 41*). Thus, cytokine produced from infection influences the course of disease (*42, 43*). An early protective Th1 response favoring IFN-γ production in the presence of IL-12 would be beneficial for a host to prevent virus infection. Immunization of mice using E1/sE2_F442NYT_-mRNA-LNP vaccine preparation, and resistance to surrogate recombinant VV E1-E2-NS2_aa134-966_ challenge infection supports the potential use of mutated E2 in a HCV vaccine. In chronically HCV infected patients, T-cells are inefficient and defective in producing IFN-γ (*44*). We have shown that interfering with HCV E2/CD81 mediated immune regulation may lead to enhanced immunogenicity for effective vaccine associated protection against infection. Reducing E2/CD81 binding results in an enhanced Th1 response and will be appropriate for selection of mRNA representing modified E2 protein as a candidate for HCV vaccine antigen. The use of modified E2 mRNA-LNP vaccine will be the most flexible approach to facilitate multigenotype based HCV vaccine.

HCV induces IL-10 in monocyte-derived DCs and inhibits DC-mediated antigen-specific T-cell activation. IL-12 activates NK cells, and induces differentiation of naïve CD4^+^T lymphocytes to become IFN-γ-producing Th1 effectors in cell-mediated immune responses to intracellular pathogens (*45*). IFN-γ priming is a positive feedback mechanism for more robust IL-12 production in certain immune responses. Th1 lymphocytes are initially activated by antigen presenting cell derived IL-12 upon pathogen infection.

Individuals who spontaneously clear HCV display broad CD4^+^T-cell responses, stronger T-cell proliferation; and higher IL-2, IFN-γ, and TNF-α production than individuals who develop chronic infection dominated by Tregs and IL-10 production (*46-51*). Strong HCV-specific CD8^+^T-cell responses are generated, and HCV-specific memory T cells persist after recovery from infection (*19, 52-54*).

An effective vaccine is a major unmet need to achieve global elimination of HCV, and there is potential for HCV vaccine development (*6, 55-57*). A recent report (*12*) suggested that an HCV T-cell based vaccine regimen encoding virus non-structural NS proteins generated specific T-cell responses, and lowered the peak HCV RNA level, but did not prevent chronic HCV infection. We have shown that cells endogenously expressing HCV proteins have perturbed HLA-DR cell surface expression (*26*) associated with the negative regulation of HLA-DR promoters at the transcriptional level. HCV E2 induces immune regulatory cytokine IL-10 and CD163 protein expression in primary macrophages (*13*), and inhibits the function of T-, B- and NK- cells through binding with CD81 (*14-16, 58*). HCV-specific CD4^+^T-cell immune deficiency may act as a primary cause of CD8^+^T-cell functional exhaustion (*18, 19*). The result from our study provides important information for the use of a modified HCV E2 antigen in a candidate vaccine and in uncovering mRNA as an ideal vaccine platform for induction of robust protective immune response with the potential to provide an effective vaccine efficacy. In this study, the use of modified E2 combined with HCV E1 on a mRNA-LNP platform induced improved immunity in a mouse model. The results from our proposed study will considerably advance highly effective vaccine development against persistent hepatotropic HCV cross-genotype specific infection using the mRNA-LNP vaccine approach and evaluate the use of modified E2 mRNA-LNP as well for a therapeutic vaccine.

## MATERIALS AND METHODS

### Cells

Human embryonic kidney (HEK) 293T, human monocytic derived THP-1 cell line, human monocyte derived primary macrophages, and murine primary peritoneal macrophages were used. HEK293T cells were maintained in DMEM containing penicillin/streptomycin, 10% fetal calf serum (FCS), 1% L-glutamine and 1% non-essential amino acids. THP-1 cells were maintained in RPMI medium containing 1% penicillin-streptomycin supplemented with 10% FCS, 1% L-glutamine, 25 mM HEPES and 12.5 nM PMA. Primary macrophages were maintained in RPMI containing penicillin/streptomycin supplemented with 10% FCS and 1% L-glutamine.

### Site-specific mutagenesis of E2

HCV (H77c genotype 1a) E2 sequence corresponding to 383-661 amino acids cloned into the pcDNA 3.1 vector (gift from Heidi E. Drummer) was isolated from transformed *E. coli* and used as a template for generation of E2 mutants by a QuikChange Lightning Site Directed Mutagenesis Kit (Agilent Technology) following the supplier’s procedure. We chose to express E2 as C-terminal truncations in mammalian cells, since soluble and correctly folded E2 can be obtained by deleting the transmembrane domain as previously reported (*59*).

sE2 single mutant (L427Y) was generated by replacing a non-polar leucine (L) amino acid residue at position 427 to a polar amino acid tyrosine (L427Y). sE2 double mutant (F442NYT) was generated on the same E2 harboring single mutant sequence by replacing two amino acids. One replacement was done at position 442 with a non-polar amino acid phenyl alanine (F) to a polar amino acid asparagine (F442N) and another mutation at position 444 of a polar amino acid glutamine (Q) with a polar amino acid threonine (Q444T). The alterations at amino acid positions 442 and 444 (FYQ→NYT) included N-linked glycosylation site based on earlier publication (*25*) and observed a significant reduction in CD81 binding. Similar observations were made by alanine replacement at amino acid position 442 and observed inhibition of CD81 binding, as well as loss of a neutralizing epitope (*60*). Single mutant (L427Y), or double mutant (F442NYT) are referred in text as sE2_L427Y_, and sE2_F442NQT_, respectively. L427Y forward primer: 5’-gcacatcaatagcacggcct ataactgcaatgaaagccttaa-3’, reverse primer: 5’-ttaaggctttcattgcagttataggccgtgctattgatgtgc-3’; F442NYT forward primer: 5’-accggctggttagcagggctcaa ctatacgcacaaat-3’, reverse primer: 5’-atttgtgcgtatagttgagccctgctaaccagccggt-3’ used for site specific mutagenesis. The constructs were confirmed by plasmid isolation and DNA sequencing.

### Cell transfection, protein expression and purification

HEK 293T cells (5×10^5^) were seeded on a 6 well plate and incubated at 37^0^C in a 5% CO_2_ overnight. The plasmid DNA isolated from clones were used for transfection using Lipofectamine™ 3000 Transfection Reagent (Thermo Fisher) following supplier’s instructions. Culture supernatant was harvested from plasmid transfected cells after 48 h. The culture supernatant was passed through a 0.45 µm filter (Nalgene) and stored at - 20^0^C until purification. The recombinant proteins containing a hexa-histidine tag was purified by passing through the nickel-NTA-affinity Column (HisPurTM Ni-NTA Spin Columns, Thermo Fisher) following supplier’s instruction. The purified protein was verified by Coomassie Blue staining and western blot analysis using anti-His antibody (Sino Biological).

### E2-CD81 binding assay

Recombinant human CD81 protein (R &D Systems) was coated (4µg/ml) on ELISA plate (Thermo Fisher Scientific) and incubated overnight at 4^0^C. Next day, the plate was washed with PBS containing 0.05%Tween-20. The wells were blocked with buffer containing 3% BSA. The plate was washed with wash buffer and 100 or 200µl HEK293T transfected culture supernatant or 1.25, 2.5 and 5.0 µg/ml of purified protein (in triplicates) were added to the wells and incubated at room temperature for 2 h. The plate was washed with buffer and 100 µl HRP-conjugated anti-his antibody (Sino Biological) was added to each well and incubated at room temperature for 1 h. The plate was washed, TMB substrate added, and the reaction was stopped for reading at 450 nm using a microplate reader.

### Stimulation of peripheral blood mononuclear cells (PBMC)

Fresh blood was collected from healthy donors in a sodium heparin vial and PBMCs were isolated by Ficoll-paque™ PLUS (GE healthcare) density gradient centrifugation. Cells (1×10^6^)/well were seeded on plate and incubated overnight at 37^0^C in a CO_2_ incubator. After 24 hrs, non-adherent cells were removed by gently washing twice, and adherent cells were incubated a further 5 days for maturation of monocytes to macrophages. On day 5, macrophages were stimulated with 2.5 µg/ml sE2 or sE2_L427Y_ for 24 h at 37^0^C. Culture supernatants were collected in fresh sterile tubes, and centrifuged at 3,000 rpm for 10 min at 18-20^0^C. Supernatants were transferred to a new sterile tube and stored at -20^0^C. Adherent cells were lysed with 100 µl 2x SDS loading buffer and stored at -80^0^C.

### Cytokine quantification

Secretory cytokines present in the cell culture supernatant of macrophages exposed to HCV sE2, sE2_L427Y,_ or sE2_F442NYT_ after 24 h were quantified using commercially available paired antibodies for IL-10, IL-12, IL-6, IFN-γ and TNF-α cytokine kit (Sino Biological). IL-4 cytokine was quantified using a kit (Thermo Fisher) following supplier’s protocol. Mouse cytokine levels in vaccinated animals (IL-4, IL-10, IL-12, and IFN-γ) was measured by ELISA following the supplier’s protocol (Invitrogen or R&D Systems).

### Macrophage/T-cell co-culture and Th1 and Th2 marker analyses

PBMCs isolated from healthy donors (n=6) were used for CD4^+^T cell isolation kit (Miltenyi Biotec). CD4^+^T cells were treated with recombinant IL-2. Matured monocyte derived macrophages were (1×10^5^) were seeded per well in a 48 well plate, and stimulated with 2.5 µg/ml sE2 or sE2_F442NYT_ for 24 hrs. Live T cells (2.5×10^4^) and macrophages (1×10^5^) were co-cultured for 4 days. On day 5, cells were harvested and stained with CD4-FITC, CXCR3-PE and CCR4-APC antibodies (BioLegend) for FACS analysis.

### CD4^+^T cell proliferation

PBMCs were isolated from healthy donors (n=3), and CD4^+^ T cells were isolated using human CD4^+^T cell isolation kit (Miltenyi Biotec). Purified CD4^+^T cells were washed and stained with 3 µM CFSE (Invitrogen). Cells were washed, activated with 5µg/ml human anti-CD3 antibody and 1µg/ml human anti-CD28 antibody (BioLegend), and incubated in the presence or absence of 2.5µg/ml sE2 or sE2_F442NQT_ for 3 days. Cells were processed for Flow cytometry.

### T-cell activation marker analysis

Isolated CD4^+^T cells from healthy donors were incubated with sE2 or sE2_F442NYT_ in the presence of 5µg/ml human anti-CD3 antibody and 1µg/ml human anti-CD28 antibody (BioLegend) for 2 hrs. Cells were harvested and stained with CD69-FITC and CD25-PE (BioLegend) for FACS analysis.

### Generation of the mRNA encoding the luciferase, and HCV protein

Codon optimized luciferase and HCV E2 specific sequence were cloned into an mRNA production plasmid (optimized 3’ and 5’ UTR and containing a polyA tail), *in vitro* transcribed in the presence of N1-methylpseudouridine modified nucleoside (N1mΨ, modified), co-transcriptionally capped using the CleanCap™ technology (TriLink) and cellulose purified (*61, 62*) to remove double stranded RNA. Purified mRNA was ethanol precipitated, washed, re-suspended in nuclease-free water, subjected to quality control (electrophoresis, dot blot, endotoxin determination using the colorimetric LAL assay and transfection into human DCs), and stored at -20°C until use.

### Generation of mRNA-LNP vaccines

mRNA loaded LNPs were formulated using a total lipid concentration of 40mM as previously described (*62*). The ethanolic lipid mixture comprising ionizable cationic lipid, phosphatidylcholine, cholesterol and polyethylene glycol-lipid was rapidly mixed with an aqueous solution containing cellulose-purified N1-mΨ *in vitro* transcribed mRNAs. The LNP formulation used in this study is proprietary to Acuitas Therapeutics (the composition is described in US patent US10,221,127). RNA-loaded particles were characterized for their size, surface charge, encapsulation efficiency and endotoxin content and subsequently stored at -80°C at an RNA concentration of 1 μg/μl (in the case of loaded particles) and total lipid concentration of 30 μg/μl (both loaded and empty particles). The mean hydrodynamic diameter of mRNA-LNPs was ∼80 nm with a polydispersity index of 0.02-0.06 and an encapsulation efficiency of ∼95%. Two or three batches from each mRNA-LNP formulations were used in these studies.

### mRNA-LNP vaccine immunization and protection against recombinant VV/HCV_1-967_

Balb/c mice (Jackson Lab) were divided into 3 groups (5 mice per group) and immunized intramuscularly in the hind leg with mRNA-LNP candidate vaccine as E1/sE2 or E1/sE2_F442NTY_ (10μg of vaccine antigen) twice at 2-week intervals. Test bleeds (3 days before immunization and 3 days before challenge infection) from mice were analyzed. Immunized mice were challenged intraperitoneally with live recombinant VV/HCV_1-967_ (*63*), and sacrificed 4 days after challenge infection and collected the ovaries for further analysis. Ovaries were homogenized with a motor operated disposable sterile homogenizer, centrifuged for clarity and used for vaccinia virus recovery by plaque assay on BSC-40 cells. Plaques were stained with crystal violet and counted.

### Statistical analysis

Student’s two-tailed paired t-test or unpaired t-test was used to determine the difference within the groups studied. One-way ANNOVA was used to compare between the groups. All the data are expressed as Mean ± SE (standard error of the mean), and p < 0.05 was considered statistically significant.

## Acknowledgements

We thank Michael Houghton, University of Alberta, Canada, for providing recombinant VV_1-967_; and Heidi Drummer, Burnet Institute, Australia, for providing the HCV sE2 clone. We acknowledge the technical help of Ms. Jenni Franey from Comparative Medicine, Saint Louis University for technical help in vaccinia challenge experiments.

## Author contributions

Conceptualization: RR conceptualized the research plan of the study and DW conceptualized the use of mRNA-LNP vaccine approach.

Methodology: VM, TP, KM, AMG, ER

Investigation: VM, TP, KM, AMG, ER, DW, RR

Funding acquisition: RR, DW

Writing- VM, TP, KM, AMG, ER, RR

Review and Editing: RR and DW

## Data and materials availability

Data associated with this study are presented in this paper.

